# Effects of seasonality, fertilization and species on the chlorophyll *a* fluorescence as related with photosynthesis and leguminous tree growth during Amazonian forest restoration

**DOI:** 10.1101/2020.11.17.386342

**Authors:** Roberto Kirmayr Jaquetti, José Francisco de Carvalho Gonçalves, Henrique Eduardo Mendonça Nascimento, Karen Cristina Pires da Costa, Jair Max Fortunato Maia, Flávia Camila Schimpl

## Abstract

The ability of species to adjust their light energy uptake is determined during plant establishment and development. Changes in resource availability may impact energy fluxes and photosynthesis. General and specific variations in chlorophyll *a* fluorescence under high vs. low water and nutrient conditions have been studied. N_2_-fixing leguminous trees, which are commonly used in tropical forest restoration, seem to be very well adapted to degraded ecosystems. To understand the effects of biological nitrogen fixation on Chl *a* fluorescence variables, three of the six Fabaceae species selected for this study were N_2_-fixing species. Additionally, the correlation among Chl *a* fluorescence and growth, photosynthesis and nutrient levels was evaluated. A 24-month forest restoration experiment was established, and data on dark-adapted Chl *a* fluorescence, photosynthesis, diameter growth and foliar nutrients were collected. Multivariate analysis was performed to detect the effects of seasonality and fertilization. Under high water- and nutrient-availability conditions, plants exhibited enhanced performance index values that were correlated with electron transport fluxes. Under drought and nutrient-poor conditions, most species exhibited increased energy dissipation as a method of photoprotection. Great interspecific variation was found; therefore, species-specific responses to the test conditions should be considered in future studies. N_2_-fixing species showed increased performance index and maximum fluorescence values, indicating their ability to colonize high-light environments. Negative correlations were found between photosynthesis and trapped fluxes and between diameter growth and initial fluorescence. Electron transport fluxes were positively correlated with growth. Given the different responses identified among species, Chl *a* fluorescence is considered a cost-effective technique to screen for seasonality, nutrient and N_2_-fixing species effects and should be considered for use during forest restoration. Finally, including N_2_-fixing species and multiple fertilization treatments in related studies may greatly facilitate the restoration of biogeochemical cycles in the tropics.

## Introduction

Plant species use light from the sun as their primary source of energy for photosynthesis. The excessive light energy in degraded tropical forest ecosystems (above 3000 μmol m^−2^ s^−1^) may cause photoinhibition in poorly adapted species, affecting their photosynthesis and growth (1,2). Some species may exhibit enhanced light uptake and plastic responses under high-light environments, and these species are considered fundamental for the establishment of forest restoration vegetation (3,4). N_2_-fixing tree species may enhance light uptake, particularly in nutrient-limited soils (5,6).

Dark-adapted chlorophyll *a* fluorescence (ChF) measurements based on dissipative (*DI*), absorbed (*ABS*), trapped (*TR*) and transport (*ET*) energy fluxes are considered effective indicators of the effects of stress on photosynthetic performance (7,8). The OJIP curve (shape) has also been applied in the early detection of different abiotic stresses (9). Various effects on the quantum yield of PSII photochemistry (*F*_V_/*F*_M_) and the performance index have been reported under drought depending on the water deficit severity and on the tested species (10).

As drought progresses, plants may downregulate electron transport and increase dissipation fluxes through the inactivation of PSII *RC*s (11,12). Degraded areas with multiple nutrient deficiencies and decreased soil organic matter in the Amazon Basin can influence plant photochemical activities, thus restricting photosynthesis and plant growth (13,14). Increased drought resistance after fertilization has been reported to be due to the enhanced quantum yield and photosynthesis (15,16). Studies on tropical environments have demonstrated the synergistic effects of multiple-nutrient fertilization on biomass growth during forest restoration with leguminous trees (17,18). However, only a few studies have evaluated the effects of combined drought and nutrient stresses on growth and photosynthetic perforance.

There has been increased interest in ChF responses to environmental cues and their correlation to photosynthesis and growth traits (19–21). ChF has been demonstrated to be an effective technique to detect different environmental cues, although the large number of variables based on *ABS*, reaction centers (*RC*s) and cross-sections (*CS*s) may confuse researchers (22). Multivariate analyses such as principal component analyses (PCAs) may reduce the noise associated with large ChF datasets, allowing researchers to understand how different species and ecological groups adjust energy fluxes (23). Likewise, in addition to being less expensive than photosynthesis measurements, ChF can also detect effects such as seasonality, which are harder to notice during growth analysis under field conditions.

Understanding ChF responses in ecological restoration species may increase forest restoration success. While a substantial number of ChF studies exist, some knowledge gaps still remain. For instance, no ecophysiological studies on important *Clitoria* or *Cenostigma* species have been published. Studies of the effects of drought on ChF traits in tropical leguminous trees are also lacking. Moreover, few studies have demonstrated the correlations between ChF variables and growth, photosynthesis and nutrient traits. Therefore, three main hypotheses were tested: 1. Leguminous trees adjust their photochemical energy fluxes under different water and nutrient availability conditions; 2. Evolutionary N_2_-fixing species will exhibit enhanced transport fluxes and plant performance regardless of nutritional status; and 3. ChF variables are correlated with growth, photosynthesis and nutrient traits.

## Materials and methods

### Experimental trial, species and treatments

Substantial variability in precipitation, temperature and soil properties can be found across Amazonian ecosystems. Central Amazonia is characterized by low natural soil fertility and high irradiance, temperatures, and monthly precipitation. Without delving into a geopolitical and scientific discussion, large- and small-scale degradation is widespread throughout the basin. The experimental trial was established in a typical homogeneous area of the Forest Restoration Program of the Balbina Hydropower Dam in Amazonas state, Brazil. Located two hours by car from Manaus, the 3 ha area was degraded and abandoned 30 years ago by the complete removal without burning of the nonflooded and dense *terra firme* forest. The area is surrounded by a natural nonflooded and dense *terra firme* forest. Despite maintaining its good physical characteristics of the soil, nutrient deficiencies for nitrogen (N) (0.16 g kg^−1^), phosphorus (P) (0.14 mg dm^−3^), calcium (Ca) and magnesium (Mg) (0.3 cmol_c_ dm^−3^), were found. Natural regeneration was not observed before intervention supporting the planting seedlings choice. Nine blocks (n = 9) measuring 6 m x 72 m (432 m^2^ area) each were placed across the area containing all 12 treatments in a combination of six tree legumes species under low- (unfertilized) and high-nutrient (fertilized) tratments (Fig 1). The 108 studied plants were selected from a total of 432 individuals planted. The high-nutrient treatment received four applications of macro- and micronutrients in the beginning of dry and wet season, while the low-nutrient treatment received no fertilization throughout the 24-month experiment.

**Fig 1.**
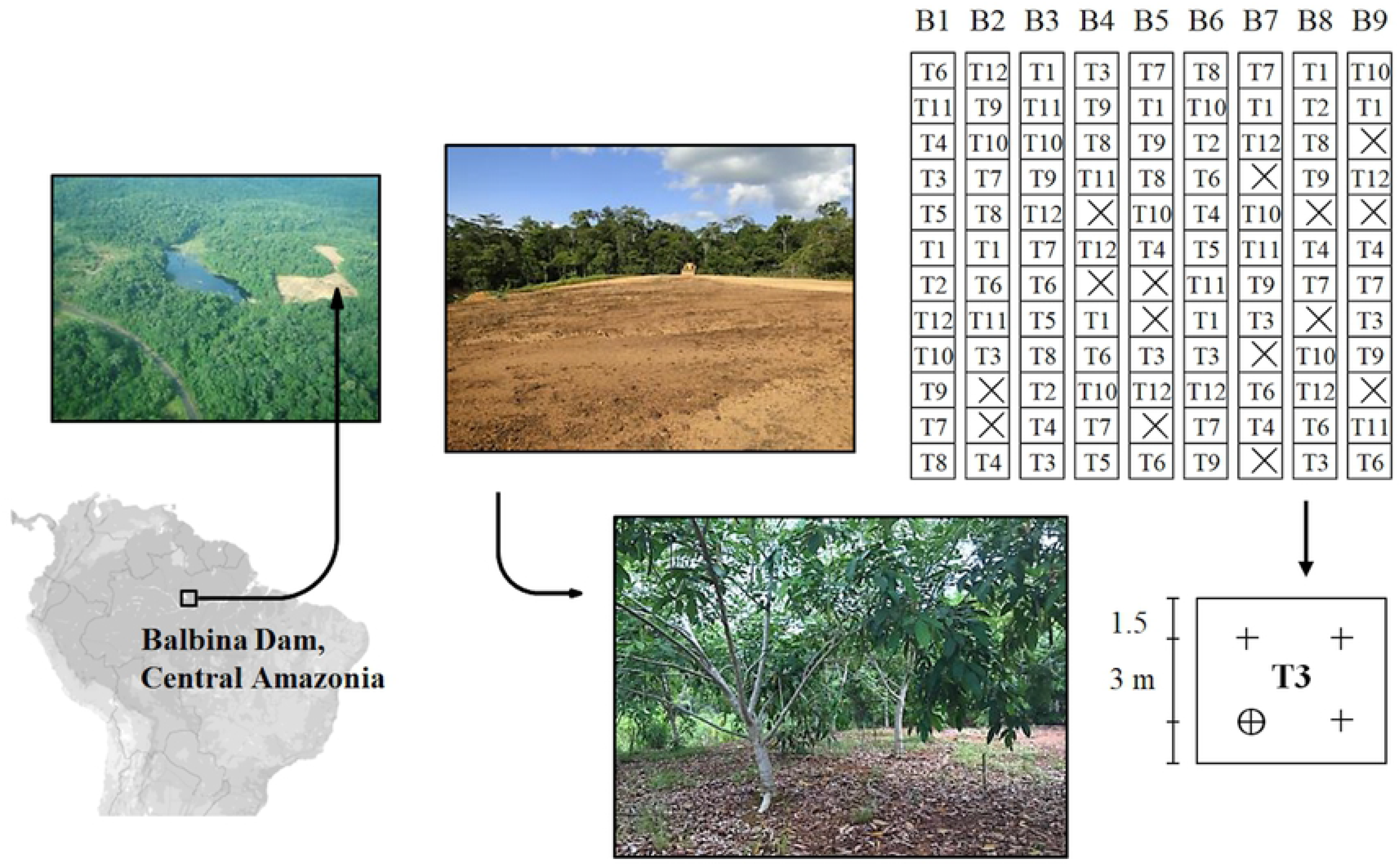
Graphical depiction with the location of the *“Alalau”* degraded area, the experimental design, and the apparent conditions before intervention and 2 years after the tropical forest restoration. Experimental design with 12 treatments (fertilization x species) randomly allocated in nine blocks (n = 9). Lost parcels (n) due to the mortality of four replicates are represented by the crossed squares (☒). T1 and T7, *C. tocantinum*; T2 and T8, *S. reticulata*; T3 and T9, *D. odorata*, T4 and T10, *C. fairchildiana*; T5 and T11, *I. edulis*; T6 and T12, *Acacia* sp.; T1 to T6, low-nutrient treatments; and T7 to T12, high-nutrient. For example, the four replicates of T3 treatment (low-nutrient x *D. odorata*) on block eight, are defined with the cross symbol (╋) in the enlarged parcel, and the circulate-cross (⨁) illustrate the selected plant for data sampling. The map was accomplished with scribblemaps.com free version.

Under normal conditions precipitation around the area is well distribuited with less rainy months (around 150 mm month^−1^) from June to November. During the experimental course the strong 2015/16 El Niño-Southern Oscillation (ENSO) event caused a 60-day period with no precipitation when first measurements were performed (dry season). Second measurements were accomplished 6 months later (rainy season), after recovery. The inclusion of inoculated N_2_-fixing species was expected to improve resource use efficiency by lowering the necessary external inputs and the time required to rebuild C stocks (24). The six species selected from Fabaceae for planting were three non-N_2_-fixing: *Cenostigma tocantinum* Ducke, *Senna reticulata* (Willd.) H.S. Irwin & Barneby, *Dipteryx odorata* (Aubl.) Willd.; and three N_2_-fixing: *Clitoria fairchildiana* R.A. Howard., *Inga edulis* Mart.—and one alien *Acacia* sp. The alien *Acacia* sp. is frequently planted as capable of growing even in highly degraded areas.

The relative growth rate of the diameter (RGR_D_) of each tree, hereafter the diameter growth, was calculated 24 months after plantating (17). Detailed information on the species used, fertilization methods, experimental trials and monthly precipitation during the experimental period can be found in (25).

### Photosynthesis measurements

Photosynthesis was measured between 8:00 and 11:30 h in nine selected plants per treatment in the dry and wet seasons. Healthy, sun-exposed and completely expanded leaves on the east side of the plants were selected from the middle third of each plant. The net photosynthetic rate (*P*_n_), hereafter referred to as photosynthesis, was measured using a portable photosynthesis system (Li-6400, Li-Cor Inc., Lincoln, NE, USA) as described by (26). Each measurement was performed at photosynthetic photon flux densities of 0 and 2000 μmol m^−2^ s^−1^ with the foliar chamber adjusted to a CO_2_ concentration, temperature and water vapor concentration of 400 ± 4 μmol mol^−1^, 31 °C ± 1 °C and 21 ± 1 mmol mol^−1^, respectively.

### Chlorophyll *a* fluorescence

The dark-adapted ChF measurements were performed with a portable fluorometer (Handy PEA, MK2-9600-Hansatech, Norfolk, UK) on the same individuals and leaves used for the photosynthesis measurements between 8:30 and 11:00 h. The selected leaves were dark-adapted for a period of 30 min and then exposed to a 1 s excitation pulse of intensely saturating light (3000 μmol m^−2^ s^−1^) at a wavelength of 650 nm. Fast fluorescence transients were calculated based on the so-called “JIP-test” (27). The performance index on an absorption basis (*PI*_ABS_), hereafter the performance index, which combines *DI*, *TR* and *ET* fluxes, was calculated as described by (28). The units and formulae used are provided in Table S2.

### Foliar nutrient concentration

The macro- and micronutrient concentrations of the leaves were determined in each treatment after the leaf samples were oven-dried at 65 °C and ground. The N concentration (mass basis) was determined with a 2400 Series II CHNS/O Organic Elemental Analyzer (PerkinElmer Inc., Waltham, MA, USA). The P concentration was determined by spectrophotometry at 725 nm. The potassium (K), Ca and Mg, iron (Fe) and zinc (Zn) concentrations were determined using atomic absorption spectrophotometry (PerkinElmer 1100B, Inc., Waltham, MA, USA) according to Jaquetti et al. (17).

### Data analysis

A complete randomized block experimental design was used. The interrelationships among ChF variables were assessed using the PCA ordination method, which reduces the dimensionality of the original data (29). PCA was performed to evaluate the effects of the seasonality and fertilization treatments. All variables were standardized by the maximum relatedness (30) prior to analysis. Product-moment correlations were used to assess the influences of seasonality (dry and wet seasons) and fertilization (fertilized and unfertilized treatments) on the ordination axes and each original variable. At a probability level of *P* < 0.05, pairwise t-tests were performed to evaluate the significance of the seasonality and fertilization effects. The analyses were run initially with 21 ChF variables and were run again with the 11 most responsive after removing similar and nonresponsive variables (S1 Table). The effects of seasonality and fertilization on the performance index and the energy dissipation flux per active PSII (*DI*_0_/*RC*), hereafter the energy dissipation, were compared using repeated measure two-way ANOVA with seasonality (dry and wet) as the repeated measure and species and fertilization treatments as factors. The relationships among ChF variables and functional traits were tested using nonparametric Spearman pairwise correlation analysis in the fertilized plants and during the wet season. The PCA was performed with PAST-UiO 3.0 (Hammer and Harper, Oslo, NO), and the inferential tests were performed with STATISTICA 12.0 (TIBCO Software Inc., CA, USA).

## Results

### Seasonal effects in the high- and low-nutrient treatments

Corroborating the first hypothesis, plants in the high-nutrient treatment adjust energy fluxes to seasonality. Significant differences separating dry and wet seasons were found in PCA axis 1 (t = −3.12, *p* < 0.01) (S1 Table). Most individuals enhanced energy dissipation fluxes and apparent antenna size of an active PSII (*ABS*/*RC*), hereafter antenna size, in the dry season (see S2 Table for the product-moment correlations). Enhanced efficiency of electron transport after Q_A_^−^ (*ET*_0_/*TR*_0_), hereafter electron transport, and performance index were found in the wet season (Fig 2). Low-nutrient plants adjust poorly to seasonal effects whit no clear separation between dry and wet seasons. Slight differences were found in PCA axis 1 (t = −2.05, *p* < 0.05) (S1 Table and S1 Fig), suggesting that multiple nutrient deficiencies weakened the leguminous trees ChF adjustments to seasonal effects.

**Fig 2.**
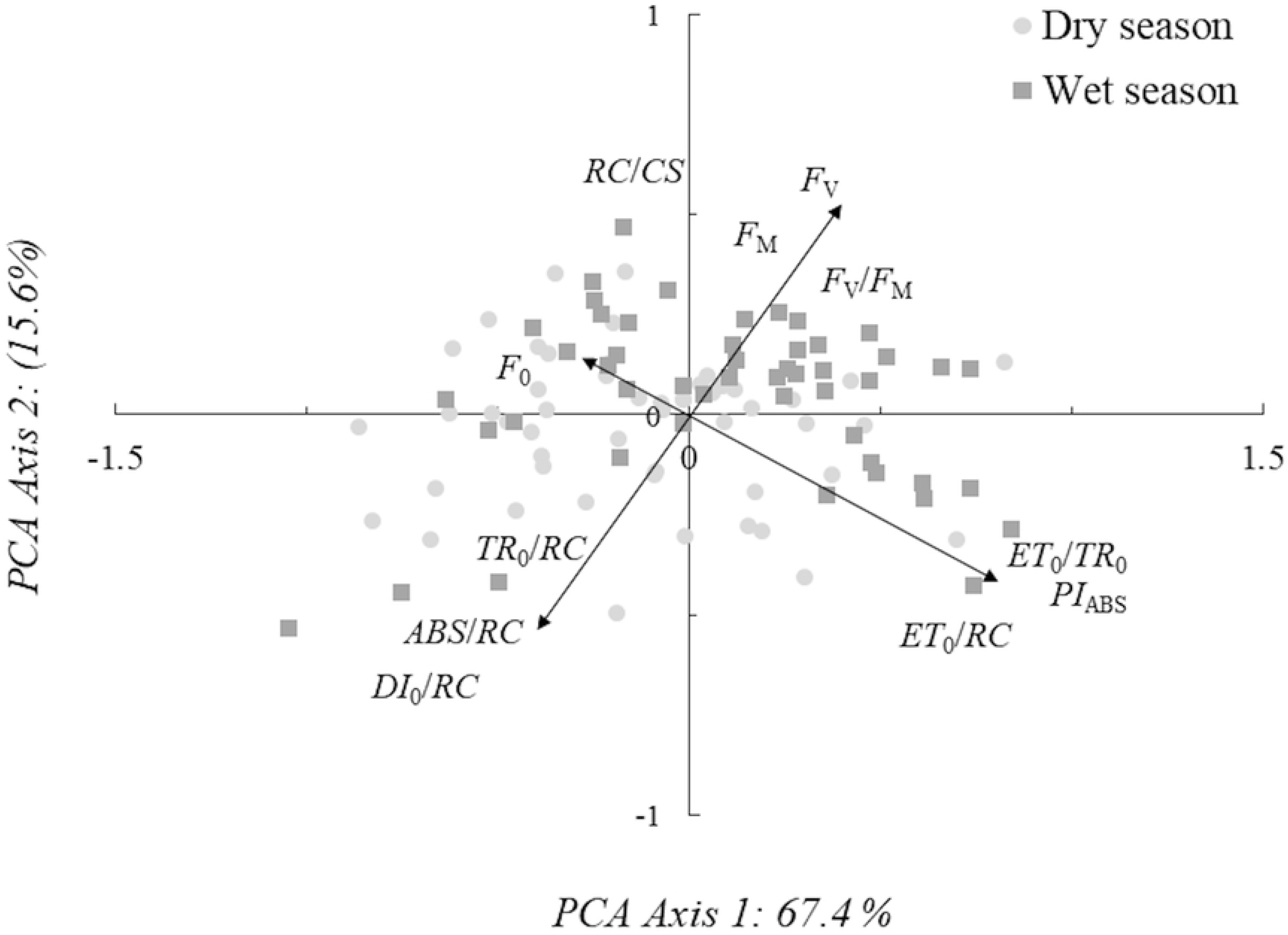
PCA ordination and loadings graphic of seasonal effects in the high-nutrient treatment. *PI*_ABS_, performance index; *ET*_0_/*TR*_0_, efficiency of electron transport; *DI*_0_/*RC*, energy dissipation flux; *ABS*/*RC*, antenna size of an active PSII *RC*; *F*_M_, maximum fluorescence; *F*_V_, maximum variable fluorescence.

### Fertilization effects during dry and wet seasons

Most plants adjust energy fluxes to different nutrient regimes during the wet season. An evident separation between high- and low-nutrient treatments was found in PCA axis 1 (t = −3.14, *p* < 0.01) (S1 Table). Plants in the high-nutrient treatment exhibited higher performance index values and greater electron transport, while low-nutrient plants exhibited increased energy dissipation and initial Chl *a* fluorescence (*F*_0_), hereafter the initial fluorescence (Fig 3). In the dry season, a noticeable effect for the fertilization treatment was found in PCA axis 1 (t = −3.61, *p* < 0.001). Low-nutrient plants contribute to increased energy dissipation fluxes, while high-nutrient plants improved electron transport and the electron transport flux per active PSII (*ET*_0_*/RC*) (S2 Fig). These findings suggest that the drought resistance of leguminous trees increased under the fertilization treatments. Low-nutrient *Acacia* sp. adjusts ChF variables differently with high electron transport and performance index values independent of seasonal effects (S3 Table).

**Fig 3.**
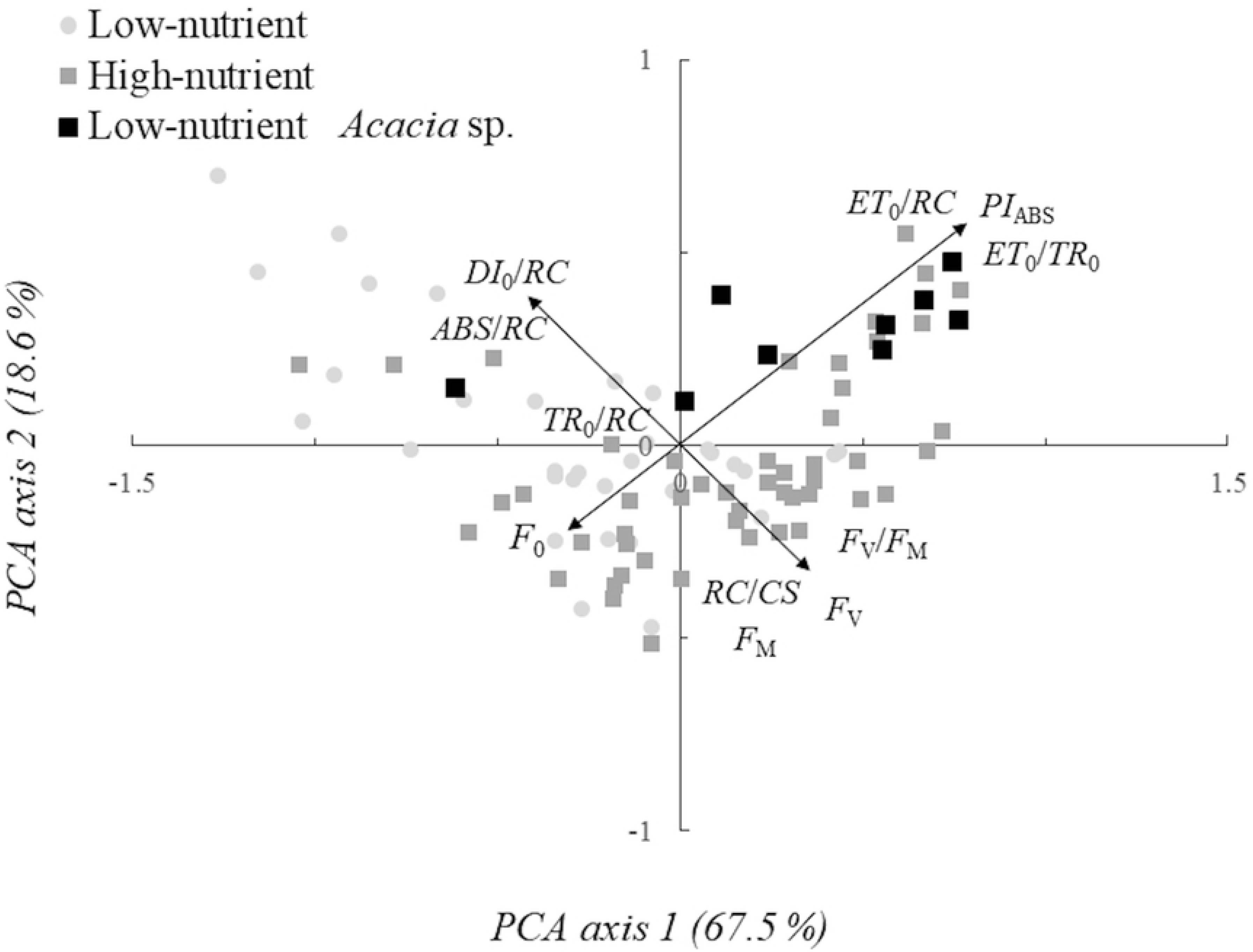
PCA ordination and loadings graphic of fertilization effects during the wet season. *PI*_ABS_, performance index on an absorption basis; *ET*_0_/*TR*_0_, efficiency of electron transport; *DI*_0_/*RC*, energy dissipation flux; *ABS*/*RC*, antenna size of an active PSII *RC*; *F*_M_, maximum fluorescence; *F*_V_, maximum variable fluorescence.

### Seasonal, fertilization and specific effects on *PI*_ABS_ and *DI*_0_/*RC*

Considering the performance index significant effects were found for the fertilization and species treatments with a significant interaction between factors. For the repeated measures differences were found between dry and wet seasons with the interaction between species and seasonal effects. No interaction was found between seasonal and fertilization effects (S4 Table). When species were compared significant differences were found between N_2_-fixing and non-N_2_-fixing species with highest performance values for *Acacia* sp. and *I. edulis* (Fig 4).

**Fig 4.**
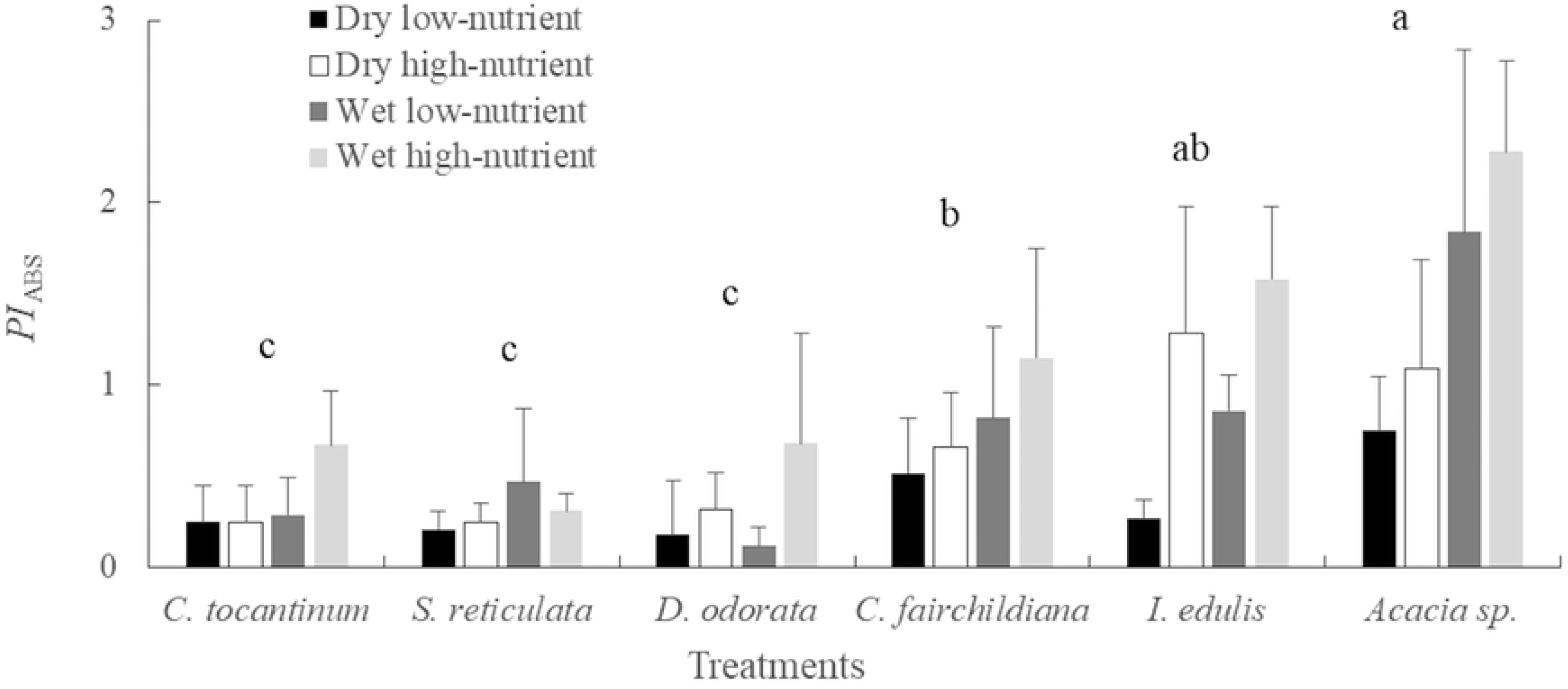
Performance index values under low vs. high water and nutrient availability (Mean ± SD).

Considering the energy dissipation significant differences were found for both fertilization and species treatments with interaction between factors. For the seasonal effects statistical differences were found among dry and wet seasons with no interactions among factors. Comparing the species, the late-successional *D. odorata* increased energy dissipation, while no differences were found among other species (Fig 5).

**Fig 5.**
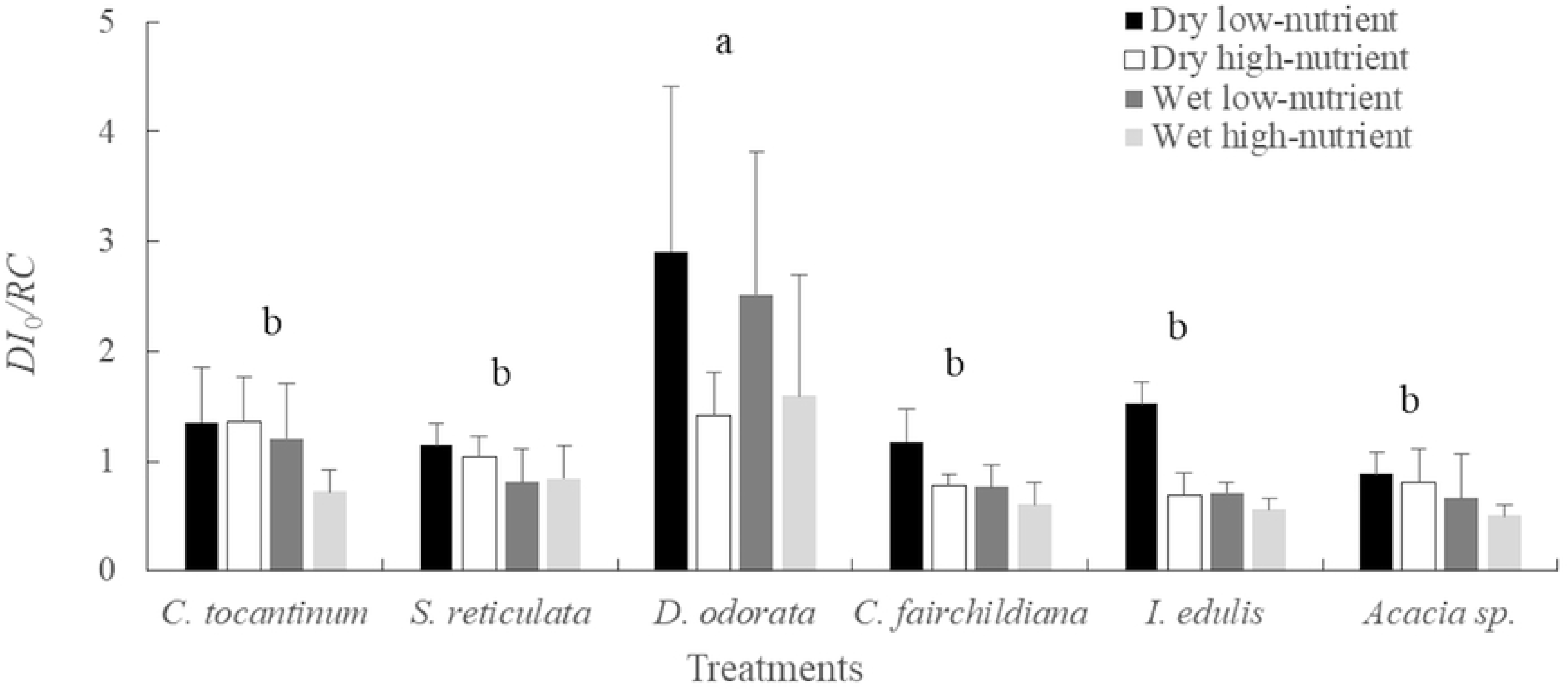
Energy dissipation flux values under low vs. high water and nutrient availability (Mean ± SD).

### Correlation of ChF with photosynthesis, foliar nutrient and plant growth

Under high water and nutrient availability, performance index was positively correlated with diameter growth (*r*_s_ = 0.65) and Zn (*r*_s_ = 0.52). Maximum trapped exciton flux of an active PSII (*TR*_0_/*RC*), hereafter the trapped flux, was negatively correlated with photosynthesis (*r*_s_ = −0.66), N (*r*_s_ = −0.58) and Zn (*r*_s_ = −0.53). Negative correlations with moderate collinearity were found between diameter growth and *F*_0_ (*r*_s_ = −0.61) and energy dissipation (*r*_s_ = −0.57) (Table 1). Electron transport was positively correlated with Zn (*r*_s_ = 0.51) and growth (*r*_s_ = 0.63) and negatively correlated with Ca (*r*_s_ = −0.58).

**Table 1:** Pairwise Spearman correlation coefficients (*r*_S_) among ChF variables and growth, photosynthesis and foliar nutrient concentrations.

|  | **RGR<sub>D</sub>** | **P<sub>n</sub>** | **N** | **P** | **K** | **Ca** | **Mg** | **Fe** | **Zn** |
| --- | --- | --- | --- | --- | --- | --- | --- | --- | --- |
| **F<sub>0</sub>** | **-0.61\*\*** | -0.35** | -0.28* | 0.00<sup>ns</sup> | 0.21<sup>ns</sup> | 0.11<sup>ns</sup> | -0.14<sup>ns</sup> | -0.28* | -0.26<sup>ns</sup> |
| **F<sub>M</sub>** | 0.31<sup>ns</sup> | -0.05<sup>ns</sup> | 0.08<sup>ns</sup> | 0.34* | -0.25<sup>ns</sup> | 0.41** | 0.02<sup>ns</sup> | -0.19<sup>ns</sup> | -0.09<sup>ns</sup> |
| **F<sub>V</sub>** | -0.13<sup>ns</sup> | 0.05<sup>ns</sup> | 0.18<sup>ns</sup> | 0.33* | -0.30* | 0.37** | 0.08<sup>ns</sup> | -0.10<sup>ns</sup> | 0.01<sup>ns</sup> |
| **F<sub>V</sub>/F<sub>M</sub>** | **0.51\*\*** | 0.29<sup>ns</sup> | 0.28* | 0.08<sup>ns</sup> | -0.27<sup>ns</sup> | 0.03<sup>ns</sup> | 0.11<sup>ns</sup> | 0.24<sup>ns</sup> | 0.22<sup>ns</sup> |
| **PI<sub>ABS</sub>** | **0.65\*\*** | 0.40* | 0.49** | -0.21<sup>ns</sup> | -0.17<sup>ns</sup> | -0.40** | -0.02<sup>ns</sup> | 0.37** | **0.52\*\*** |
| **ET<sub>0</sub>/TR<sub>0</sub>** | **0.63\*\*** | 0.38* | 0.46** | -0.32* | -0.17<sup>ns</sup> | **-0.50\*\*** | -0.08<sup>ns</sup> | 0.33* | **0.51\*\*** |
| **TR<sub>0</sub>/RC** | -0.44** | **-0.66\*\*** | **-0.58\*\*** | -0.03<sup>ns</sup> | 0.09<sup>ns</sup> | -0.41** | 0.08<sup>ns</sup> | -0.26<sup>ns</sup> | **-0.53\*\*** |
| **DI<sub>0</sub>/RC** | **-0.57\*\*** | -0.44** | -0.42** | -0.04<sup>ns</sup> | 0.23<sup>ns</sup> | 0.11<sup>ns</sup> | -0.08<sup>ns</sup> | -0.26<sup>ns</sup> | -0.35* |

$F_0$ , initial fluorescence; $F_M$ , maximum fluorescence; $F_V$ , maximum variable fluorescence; $F_V/F_M$ , maximum quantum yield of PSII photochemistry; $PI_{ABS}$ , performance index on an absorption basis; $ET_0/TR_0$ , efficiency of electron transport; $TR_0/RC$ , maximum trapped exciton flux; $DI_0/RC$ , energy dissipation flux; RGR<sub>D</sub>, relative growth rate in diameter; $P_n$ , net photosynthetic rate; N, nitrogen, P, phosphorous, K, potassium, Ca, calcium, Mg, magnesium, Fe, iron, and Zn, zinc concentrations in leaves.
Bold values are highly significant ( $p < 0.001$ ) with high ( $r_s > 0.70$ ) and moderate ( $0.50 < r_s < 0.70$ ) collinearity between variables. \* Significance at the 0.05 level; \*\* Significance at the 0.01 level.

## Discussion

### Ecofunctional aspects of seasonal and nutrient effects on ChF

As general stress responses plants exposed to drought and nutrient deficiencies in this study exhibited enhanced energy dissipation, antenna size and initial fluorescence conformingly as energy may be lost from plants through fluorescence emissions and heat dissipation (31). With high water and nutrient availability, plant energy is mainly directed to photochemistry (32) and electron transport in plants increases, thus improving plant performance (Fig 6).

**Fig 6:**
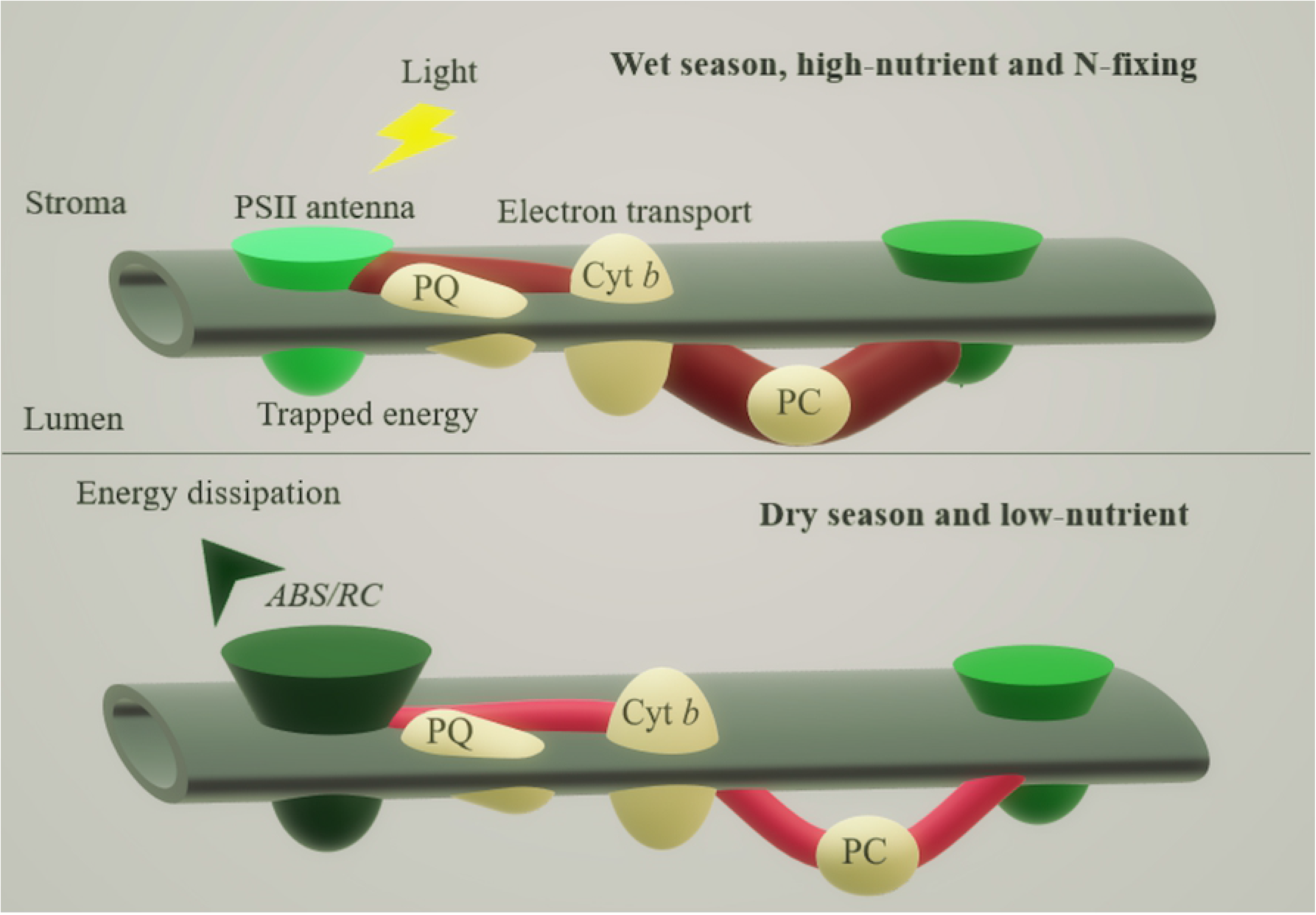
ChF adjustments in antenna size, energy dissipation and electron transport. *ABS*/*RC*, antenna size of an active PSII reaction center. The dark green means increased and light green decreased values of, dark red represent increased quantum yield and electron transport flux. Dark green and red indicates increased values of PSII antenna size and electron transport flux, and light green and red indicates decreased values.

Usually, degraded environments expose plants to excessive light energy, and early-successional species are thought to be less photoinhibited than late-successional species (33,34). Correspondingly, energy dissipation increased in the late-successional *D. odorata*, indicating photoinhibition and oxidative stress (35). Nonetheless, the late-successional *C. tocantinum* appeared to be better adapted to high-light conditions when it was fertilized. Under high-light environments, adapted species such as the N_2_-fixing *C. fairchildiana, I. edulis* and *Acacia* sp. enhance their electron transport, increasing their sink strength and light use efficiency (36). The novel finding of the positive correlation between growth and electron transport may have important applications in developing genetic improvements that will result in higher individual plant performance.

Extreme climatic events such as the ENSO may cause drought periods in previously unaffected regions of the Amazon Basin (37). Under moderate drought conditions, photosynthesis may be downregulated, which restricts CO_2_ availability, thus affecting photochemistry and electron transport (38,39). Corroborating the present findings through metabolic adjustments, drought-resistant *Acacia* species can maintain cell membrane integrity and electron transport longer than other species as the water deficit increases (40,41). The increased drought tolerance under fertilization found in three species in the present study may be related to the increased PSII RC aperture, electron transport efficiency and photoinhibition relief (42).

Plants may respond to fertilization treatments with increased plant performance and photosynthesis due to the increased nutrient availability in the soil (6,43). The decreased performance index under drought and nutrient deficiencies suggests increased photoinhibition and reduced electron transport, which negatively affect photosynthesis and growth (42).

The present study demonstrates the negative correlations between trapped energy flux and photosynthesis and between initial fluorescence and growth. Both of these findings have important potential ecological and silvicultural applications in the selection of well-adapted species for degraded environments. In addition to the present results, positive correlations were found among photosynthesis, quantum yield and variable fluorescence in *Populus* and *Miscanthus* species (44,45). Positive correlations were previously found among Fe, the performance index and electron transport (46) and between electron transport and photosynthesis (47,48). In contrast, no correlation was observed between photosynthesis and ChF variables in barley plants under drought and control treatments (12).

### Performance variables, *PI*_ABS_ and *ET*_0_/*TR*_0_

The *PI*_ABS_, which incorporates the density of PSII RCs, electron transport beyond *Q*_A_, and trapped fluxes, is widely used to study photosynthesis and the functionality of PSII and PSI under stress (1,10). The increased performance index values found under high resource availability and in N_2_-fixing species may result from changes in the PSII antenna size, trapping efficiency and electron transport (8,27). Here, multivariate analysis demonstrated a positive correlation between plant performance and electron transport adjustments. Corroborating previous findings, the performance index was a more sensitive variable for evaluating fertilization and drought effects than the PSII quantum yield (28,49).

The electron transport fluxes variables reflect the maximum electron transport between PSII and PSI and indicate changes in photosynthetic apparatus activity (50). The increased electron transport found in the well-adapted and highly productive *Acacia* sp. species may reflect improved recovery on the PSII acceptor side. The diminished electron transport reported under nutrient-poor conditions may be partially due to the overreduction of the PSII acceptor side (51).

### Stress variables, *DI*_0_/*RC*, *ABS*/*RC* and *F*_0_

Under the stressful, high-light conditions found in degraded areas, trees may increase their protective quenching activity to dissipate excessive absorbed energy The increased dissipation fluxes that occur due to the inactivation of RCs may work as strong quenchers to reduce photooxidative damage (28,53). Moreover, the increased energy dissipation indicates the inefficient reoxidation of the reduced *Q*_A_^−^ and electron transport to quinone *Q*_B_ (8,54). Corroborating previous studies, the positive correlation between energy dissipation and the initial fluorescence of stressed individuals may be associated with increased antenna size and PSII RC inactivation (11).

The negative correlation between dissipative energy and *P*_n_ suggests decreased electron transport and photochemical yield. Increased energy dissipation and antenna size have been previously reported as early indicators of the effects of high light, drought, and nutrient deficiency (9,54). Both positive and negligible effects of drought on PSII quantum yield have been previously reported; the effect depends on the nutrient status of individual plants (55,56). Our results indicate negligible interactions of seasonality and fertilization treatments effects. Moreover, in species of the *Acacia* genus, increasing water deficit has negligible effects on quantum yield, which is related to the increased protective carotenoid content (57).

### OJIP transient test

The OJIP transient curve is used for the early detection of drought and nutrient stress (58). Multiple nutrient deficiencies reduce the slope of the OJIP curves in most species due to reduced ATP production under K and P deficiencies (9,58). The reduced O-J rise found in *S. reticulata* and *C. tocantinum* reflected the impaired electron transport from pheophytin and beyond *Q*_A_ (59). The stress responses of *S. reticulata* and *D. odorata* (S3 Fig) resulted in a reduced amplitude of the thermal phase (I-P rise) due to the accumulation of reduced *Q*_A_^−^ and the decreased activity of PSII (11,32).

The sharp decrease in the maximum Chl *a* fluorescence (*F*_M_) and the increase in initial fluorescence values observed in *D. odorata* under high light suggest the inactivation of RCs and the inhibition of electron transport beyond *Q*_A_ (60). The enhanced photochemical efficiency and productivity in *Acacia* sp. were due to the increased *F*_M_ values and the absence of the P step (S3 Fig). The increased I-P rise in the well-adapted and highly productive *Acacia* sp. may be related to the enhanced rate of ferredoxin reduction and electron transport between PQ and PSI (36).

## Conclusions

The ChF technique is valuable for understanding the photochemical phase of photosynthesis and how it affects other functional traits. Nevertheless, ChF has rarely been used to assess the photosynthetic performance and adaptation ability of species used in forest restoration. As evidenced in the present study, adjustments in energy fluxes during light uptake are determinants of the growth and establishment of different species. Additionally, plants can increase energy dissipation when resources are scarce and enhance electron transport when resources are abundant. N_2_-fixing species with enhanced performance appear to be highly adapted to degraded, high-light environments. In particular, the increased electron transport fluxes in *Acacia* sp. may explain the enhanced sink strength and growth of these species in locations with multiple resource limitations. Future studies on the functional aspects of leguminous trees are recommended, especially studies on N_2_-fixing species that may facilitate the restoration of important biogeochemical cycles.

## Acknowledgments

The authors are grateful to the Balbina Hydroelectric Dam for enabling the collection of experimental data and to the Coordination for the Improvement of Higher Education Personnel (Coordenação de Aperfeiçoamento de Pessoal de Nível Superior – CAPES) and the National Council for Scientific and Technological Development (Conselho Nacional de Desenvolvimento Científico e Tecnológico – CNPq) for financial support. J.F.C. Gonçalves is a researcher with the Brazilian Council for Research and Development (CNPq).

## Supporting information

**S1 Fig. PCA ordination and loadings graphic for the seasonal effects in the low-nutrient treatment.** *PI*_ABS_, performance index on an absorption basis; *ET*_0_/*TR*_0_, efficiency of electron transport; *DI*_0_/*RC*, energy dissipation flux; *ABS*/*RC*, antenna size of an active PSII reaction center; *F*_M_, maximum fluorescence; *F*_V_, maximum variable fluorescence.

**S2 Fig. PCA ordination and loadings graphic for the fertilization effect in the dry season.** *ET*_0_/*TR*_0_, efficiency of electron transport; *ET*_0_/*RC*, electron transport flux; *DI*_0_/*RC*, energy dissipation flux; *ABS*/*RC*, antenna size of an active PSII reaction center; *F*_M_, maximum fluorescence; *F*_V_, maximum variable fluorescence.

**S3 Fig. The OJIP curve during the wet season under low (○) and high (□) nutrient conditions.**

**S1 Table. Effects of seasonality and fertilization with 21 and 11 ChF variables.**

**S2 Table. Product-moment correlations obtained through principal component analysis (PCA).** *F*_0_, initial fluorescence; *F*_M_, maximum fluorescence; *F*_V_, maximum variable fluorescence; *F*_V_/*F*_M_, maximum quantum yield of PSII photochemistry; *PI*_ABS_, performance index on an absorption basis; *ET*_0_/*TR*_0_, efficiency of electron transport; *DI*_0_/*RC*, energy dissipation flux; *ABS*/*RC*, antenna size of an active PSII reaction center; *ET*_0_/*RC*, electron transport flux; *TR*_0_/*RC*, maximum trapped exciton flux; *RC*/*CS*, density of reaction centers per cross-section.

**S3 Table. Mean ChF variables values for the six studied species under the different water and nutrient regimes.** *F*_0_, initial fluorescence; *F*_M_, maximum fluorescence; *F*_V_, maximum variable fluorescence; *F*_V_/*F*_M_, maximum quantum yield of PSII photochemistry; *PI*_ABS_, performance index on an absorption basis; *ET*_0_/*TR*_0_, efficiency of electron transport; *DI*_0_/*RC*, energy dissipation flux; *ABS*/*RC*, antenna size of an active PSII *RC*; *ET*_0_/*RC*, electron transport flux; *TR*_0_/*RC*, maximum trapped exciton flux; *RC*/*CS*, density of reaction centers per cross-section.

**S4 Table. Results of the ANOVA for the effects of seasonality, fertilization and species on *PI*_ABS_ and *DI*_0_/*RC***. *PI*_ABS_, performance index on an absorption basis; *DI*_0_/*RC*, energy dissipation flux. Degrees of freedom for species = 5, and fertilization = 1. * Significance at the 0.05 level; ** Significance at the 0.01 level.

